# Exploration of insulin amyloid polymorphism using Raman spectroscopy and imaging

**DOI:** 10.1101/782672

**Authors:** M. Ishigaki, K. Morimoto, E. Chatani, Y. Ozaki

## Abstract

We aimed to investigate insulin amyloid fibril polymorphism caused by salt effects and heating temperature, and to visualize the structural differences of the polymorphisms *in situ* using Raman imaging without labeling. The time course monitoring for amyloid formation was carried out in an acidic condition without any salts and with two species of salts (NaCl and Na_2_SO_4_) by heating at 60, 70, 80, and 90 ℃. The intensity ratio of two Raman bands at 1672 and 1657 cm^-1^ due to β-sheet and α-helix structures was revealed to be an indicator of amyloid fibril formation, and the relative proportion of the β-sheet structure was higher in the case with salts, especially at a higher temperature and with Na_2_SO_4_. In conjunction with the secondary structural changes of proteins, the S-S stretching vibrational mode of a disulfide bond (∼514 cm^-1^) and the ratio of the tyrosine doublet *R*(*I*_850_⁄*I*_826_) were also found to be markers distinguishing polymorphisms of insulin amyloid fibrils by principal component analysis (PCA). Especially, amyloid fibrils with Na_2_SO_4_ media formed the g-g-g conformation of disulfide bond at a higher rate and without any salts; on the contrary, the g-g-g conformation was partially transformed into the g-g-t conformation at higher temperatures. The different environments of the hydroxyl groups of the tyrosine residue were assumed to be caused by fibril polymorphism. Raman imaging using these marker bands also successfully visualized the two- and three-dimensional structural differences of amyloid polymorphisms. The present results indicate the potential of Raman imaging as a diagnostic tool for polymorphisms in tissues of amyloid-related diseases.

**Statement of Significance:** Our results revealed three Raman markers distinguishing amyloid fibril polymorphisms caused by salt and temperature effects; the relative proportion of protein secondary structures (α–helix and β-sheet), the ratio of tyrosine doublet, and the conformational differences of disulfide bonds. The lower values of tyrosine doublet in the case with salts were interpreted as the anions rob the hydration water from proteins which induced protein misfolding. Using these parameters, Raman images captured their higher order structural differences *in situ* without labeling. The images of hydrogen bonds strength variations due to tyrosine doublet is believed to include significant novelty. The present results imply the potential of Raman imaging for use as a diagnostic imaging tool for tissues with amyloid-induced diseases.

## Introduction

Protein aggregation and structural changes can be cited as examples of protein physical instability. Amyloid fibrils are abnormal protein aggregations, and their deposition causes diseases, such as Alzheimer’s disease, Parkinson’s disease, and type 2 diabetes (1–3). Therefore, the process of amyloid fibril formation has been investigated for decades to elucidate the mechanisms by which such diseases are caused and to prevent their onset (4, 5). Insulin is one of the proteins that easily aggregates and forms amyloid fibrils. Insulin has been used for the treatment of diabetes and studies to improve insulin stability are ongoing for its therapeutic use.

Amyloid formation proceeds through a nucleation-dependent mechanism. The nucleus, as a precursor for fibril formation, is made because of protein monomer assembly; further assembly causes elongation to form amyloid fibrils (6, 7). Insulin protein spontaneously forms amyloid fibrils without any seeds under acidic conditions with heating (8–10). The aggregation process is strongly affected by the physico-chemical environment, such as pH, temperature, ionic strength, peptide concentrations, buffer compositions, and so on. Anions are one of the most influential factors for amyloid fibril formation. The structure of amyloid fibrils and its formation speed have been reported to vary upon the addition of salts (11, 12). Raman B. et al investigated the anion effects on amyloid formation in *β*_2_-microglobulin by adding NaCl, NaI, NaClO_4_, and Na_2_SO_4_, and the order of their impact on the process was concluded to be 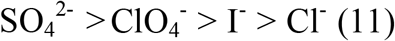, which corresponded to the Hofmeister series of anion ranking for protein precipitation (13, 14). The sodium anions were also ranked by Klement et al. as 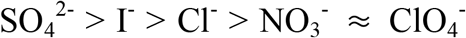 based on the aggregation propensity of the Alzheimer’s Aβ (1–40) peptide (15).

Furthermore, the different physico-chemical conditions sometimes generate polymorphs of amyloid fibrils (16, 17). Actually, brain tissues of Alzheimer’s disease showed a wide variety of morphologies of amyloid deposition (18, 19). Recent studies have clarified that polymorphs often originate from differences in microscopic structures within the protofilaments (20), although there are also some cases that arise due to differences in the number and bundle of protofilaments (21). Petkova et al. discussed possible pathways of formation of amyloid-β peptide polymorphs with variations in molecular structure (16). They concluded that the distinct morphologies were generated at the protofilament level and the seeded structures with different morphologies were self-propagating (16). It has also been reported that the morphological variants show different toxicities in cell cultures (16, 22). If the mechanisms responsible for various morphologies of amyloid fibrils (that are highly toxic) are elucidated, it might be possible to cure and prevent amyloid-related diseases. An investigation for the factors that modulate amyloid formation and generate polymorphs is very important to understand the mechanism of onset of amyloid-related diseases. The present methods to explore amyloid polymorphism are transmission electron microscope (TEM), atomic force microscope (AFM), solid-state nuclear magnetic resonance (ssNMR), and infrared (IR) spectroscopy. Very recently, cryo-EM has been applied as a very powerful tool for the structural analysis of amyloid fibrils and has revealed several polymorphs at an atomic level (23, 24). However, in terms of use as a possible tool for pathological diagnosis by investigating the structure and the distribution of amyloid polymorphisms in a non-destructive manner, *in situ*, *in vivo*, in two or three dimensions, and in real time, Raman spectroscopy and imaging are strong candidates.

Raman spectroscopy has the advantage of providing information about the molecular structure and composition *in situ* without staining (25, 27). It shows spectral patterns reflected by protein secondary structure (28, 29), environments of amino acid residues (30), and disulfide conformation (22, 31, 32). Therefore, Raman spectroscopy is a very useful tool for exploring protein structures. Ortiz et al. reported the variation of Raman spectra due to the formation of insulin amyloid fibrils in amide I, amide III, and peptide backbone regions with the increase of β-sheet components (33). Kurouski et al. analyzed insulin fibril structure by deep ultraviolet resonance Raman (DUVRR), and they found spectral patterns suggesting a characteristic disulfide conformation and local environments of tyrosine residues. They concluded that a predominance of the gauche-gauche-gauche (g-g-g) conformation remained during the aggregation process for all fibril disulfide bonds and that three out of four tyrosine amino acid residues were packed into the fibril core of insulin aggregates and another aromatic amino acid (phenylalanine) stayed in the unordered parts of insulin fibrils as indicated by the significant decrease in tyrosine band intensity (32). The distinct morphologies of amyloid fibrils are commonly visualized using TEM and AFM (16, 34–46), and the detailed structures in the order of nanometers are clearly visible. Paulite et al. succeeded in constructing tip-enhanced Raman images of nanotapes formed by β-amyloid (1–40) peptide fragments by plotting the band intensity at 1004 cm^-1^ of phenylalanine (37). All these methods are good for structural analysis at the nanometer scale, but they have some problems, for example, they take a very long time to record imaging data and are unsuitable to visualize larger-scale structures. Raman imaging, on the other hand, is a useful tool to investigate the structures in the order of micrometers, and it can obtain molecular information not only at the surface but also in the core of amyloid fibrils. Therefore, Raman imaging is a candidate of diagnostic tool to image the state of tissues with amyloid fibril deposition. However, Raman imaging of amyloid polymorphisms has hardly been attempted. So, we aimed to visualize polymorphs of insulin amyloid using Raman marker bands to distinguish polymorphisms in the order of micrometers.

In the present study, we aimed to explore the formation process of insulin amyloid fibrils by Raman spectroscopy and imaging *in situ* and investigated the polymorphisms caused by salt effects and heating temperature. Insulin samples were heated under an acidic condition without any salts, or with NaCl and with Na_2_SO_4_; these salts were selected as the anions showed different impacts on amyloid fibrillation in previous studies (13, 15). Our results revealed that Raman spectroscopy can monitor the process of amyloid fibril formation in time course by looking at the secondary structural changes of insulin protein. Perturbations, such as the addition of salt and changing the heating temperature given to insulin samples, induced polymorphisms of amyloid fibrils and their variations could be assessed by three parameters: the ratio of proteins with different secondary structures (α-helix and β-sheet), conformational differences of disulfide bonds, and the ratio of tyrosine doublets relating with the strength of hydrogen bonds of the hydroxyl group in the side chain of tyrosine residues. Raman images consistently visualize polymorphisms of amyloid fibrils caused by salt effects based on these three parameters. The present results indicate the potential of Raman imaging for use as a diagnostic imaging tool for tissues with amyloid-induced diseases.

## Materials and Methods

### Preparation of insulin sample

Recombinant human insulin (UniProtKB P01308) was purchased from Wako Pure Chemical Industries, Ltd (Japan). Insulin was diluted with 100 mM HCl to a final concentration of 100 mg/mL. Insulin solution was injected into a glass capillary (outer diameter: 1.52 mm, inner diameter: 1.13 mm, Drummond Scientific Company, USA) and both ends of the capillary were closed using a gas burner. The formation of insulin amyloid fibrils without any seeds was performed by heating under acidic conditions (8–10). The glass capillaries, after encapsulating samples, were warmed in a water bath (DTU-1CN, TAITECH, Japan) to allow amyloid fibrils formation.

In order to monitor the process of amyloid fibril formation, heating temperatures and heating times were changed. First, insulin samples were heated at 50, 60, 70, 80, or 90℃ for 20 minutes. After heating, samples were taken out from the water bath and Raman spectra were recorded at room temperature. Next, the heating time was changed every 5 minutes until the time when Raman spectra did not change any more due to protein secondary structural changes.

Furthermore, the salt effect on fibril structure was investigated by adding NaCl (100 mM) and Na_2_SO_4_ (5 mM) to the prepared insulin samples mentioned above. These two solutions were included so that the final concentrations of insulin and HCl were 100 mg/mL and 100 mM, respectively.

### Confirmation of fibril formation

In order to confirm the formation of insulin fibrils, the fluorescent pigment of thioflavin T (ThT) was used. The powder of ThT was dissolved in distilled water and mixed with a glycine sodium hydroxide buffer solution (pH=8.5). The final concentrations of ThT and buffer solution were 5 μM and 50 mM, respectively. Insulin solutions heated at five different temperatures (50, 60, 70, 80, 90 ℃) within a glass capillary were taken from the capillaries and transferred to micro tubes with a ThT solution. After stirring the mixed solutions, fluorescence measurements were performed for three samples per each heating temperature using RF-5300PC (Shimadzu, Japan). The excitation and detection wavelengths were 445 and 485 nm, respectively.

### Measurement of Raman spectra and Raman imaging

The insulin Raman spectra were recorded in the 1800-300 cm^-1^ wavenumber region with a 514 nm laser excitation wavelength, an exposure time of 10 s, and 10 scan accumulations, leading to a total exposure time of 100 s (HR-800-LWR, HORIBA, Japan). The power of the incident laser light was 46.0 mW and Raman spectra were calibrated using the peak of silicon. Raman spectra were pre-processed using background subtraction and fifth-order polynomial fitting for fluorescence background removal. All Raman spectra obtained from the samples were normalized using a standard peak at 1003 cm^-1^, due to phenylalanine. Principal component analysis (PCA) was performed to extract different spectral components between samples using the chemometrics software Unscrambler X 10.3 (Camo Analytics, Oslo, Norway).

Raman imaging data were obtained from insulin samples dissolved in three different solutions; (i) 100 mM HCl, (ii) 100 mM HCl + 10 mM NaCl, and (iii) 100 mM HCl + 5 mM Na_2_SO_4_. The insulin concentrations were unified to 100 mg/mL. The samples were heated at 90 ℃ for 30 minutes and they were measured at room temperature using an *inVia* confocal Raman microscope (Renishaw Inc., UK) system at a 532 nm excitation wavelength. The imaging data were recorded in the 1800-274 cm^-1^ wavenumber region; the exposure time was 1s, the spatial resolution was 1 μm, and the laser power was 50 mW. All images were constructed by plotting Raman band intensities or the band intensity ratios in two dimensions with same scales using the Graph-R free software.

The protein concentrations in the aqueous solution were very small compared to those of aggregations after amyloid fibrils were completely formed, and the Raman band intensities at 1003 cm^-1^ were overwhelmingly lower compared to those of aggregates. The extremely small values of the 1003 cm^-1^ band intensity inevitably enhanced the Raman band ratio defined as that divided by the 1003 cm^-1^ band intensity. In order to exclude noise components in Raman imaging due to the low concentrations of proteins, certain data points were ruled out and the ones with higher values of the band intensity at 1003 cm^-1^ compared to the threshold number characteristic for aggregation were selected. Therefore, comparison of Raman imaging is effective only for aggregations.

## Results and Discussion

Figure 1a shows the Raman spectra of human insulin solutions without any salts in the 1800-300 cm^-1^ region heated for 20 minutes at different temperatures. The top spectrum in Figure 1a was obtained from the sample without heating, and it was defined as the spectrum of the native state of insulin in the present study. A peak at 1003 cm^-1^ was due to the ring breathing mode of phenylalanine residues (28, 29); its peak height was used as an internal standard for the normalization of Raman spectra, and a band at 1031 cm^-1^ originated from the C-H in-plane bending mode of the phenylalanine residues. A peak at 642 cm^-1^ was assigned to the C-C twisting mode of the tyrosine residue, and a doublet at 850 and 826 cm^-1^ came from Fermi resonance between a ring breathing mode and the overtone of an out-of-plane ring bending vibration of tyrosine (28, 29). A band at 1207 cm^-1^ arose from a stretching mode of C-C_6_H_5_ of phenylalanine, and that at 1448 cm^-1^ arose from C-H deformation modes of proteins. Bands at 1657 cm^-1^ and 1265 cm^-1^ could be assigned to the amide I and amide III modes of proteins, respectively. These band positions at 1657 cm^-1^ of the amide I mode and at 1265 cm^-1^ of the amide III mode indicated that the insulin structure is largely α-helix; this result is consistent with the well-known fact that the native insulin has an α-helix structure (38, 39). With the increase in temperature, drastic spectral changes were observed at 80 ℃ and above. The amide I band at 1657 cm^-1^ shifted to 1672 cm^-1^ which was assigned as the protein band with the secondary structure of β-sheet (28, 29), and the band width got sharp (Figure 1b). Furthermore, a peak shift of amide III from 1265 to 1224 cm^-1^ also revealed the structural change of the protein from α-helix to β-sheet at 80℃ and above (Figure 1c) (28, 29). Notable spectral variations in Figure 1a were also observed at around 500 cm^-1^ due to disulfide bonds and the enlarged view in 600-400 cm^-1^ is shown in Figure 1d. The insulin molecule consists of two polypeptides chains named A chain and B chain with 21 and 30 amino acid residues, respectively, and they are linked by disulfide bonds to each other (40). Insulin has three disulfide bonds: two inter-chain and one intra-chain disulfide bond, and they have a crucial impact on its stability, activity, and physiological functions (22, 41). The conformations of disulfide bonds decide the tertiary and secondary structure of proteins (4, 5), and Raman spectroscopy is a useful way to observe the conformation of disulfide bonds in situ and in real time. The samples heated above 80 ℃ yielded striking peaks at around 514 cm^-1^ due to the S-S stretching mode (22, 28, 31, 32). It has been well known that Raman bands due to disulfide bonds appear in 550-500 cm^-1^ region, and the ones with different conformations exhibited at different wavenumbers; gauche-gauche-gauche (g-g-g), gauche-gauche-trans (g-g-t), and trans-gauche-trans (t-g-t) conformation give a characteristic S-S stretching band at 515, 525, and 540 cm^-1^, respectively (22, 28, 32, 42). Therefore, the results in Figure 1d indicate that amyloid fibrils mainly have a g-g-g conformation (31). The increment of band intensity due to disulfide bonds occurred with the secondary structural changes shown in the amide I and amide III regions.

**Figure 1:**
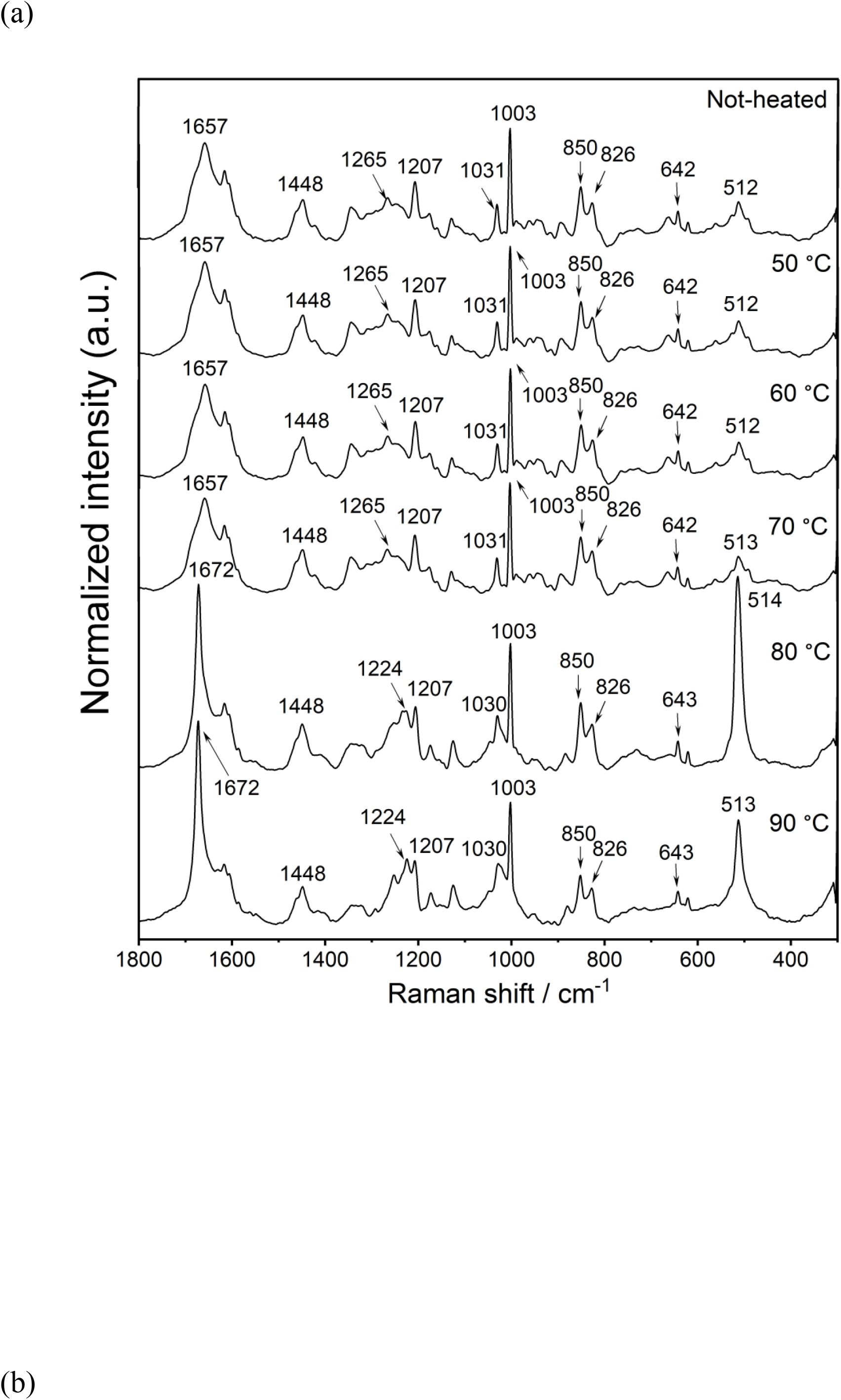
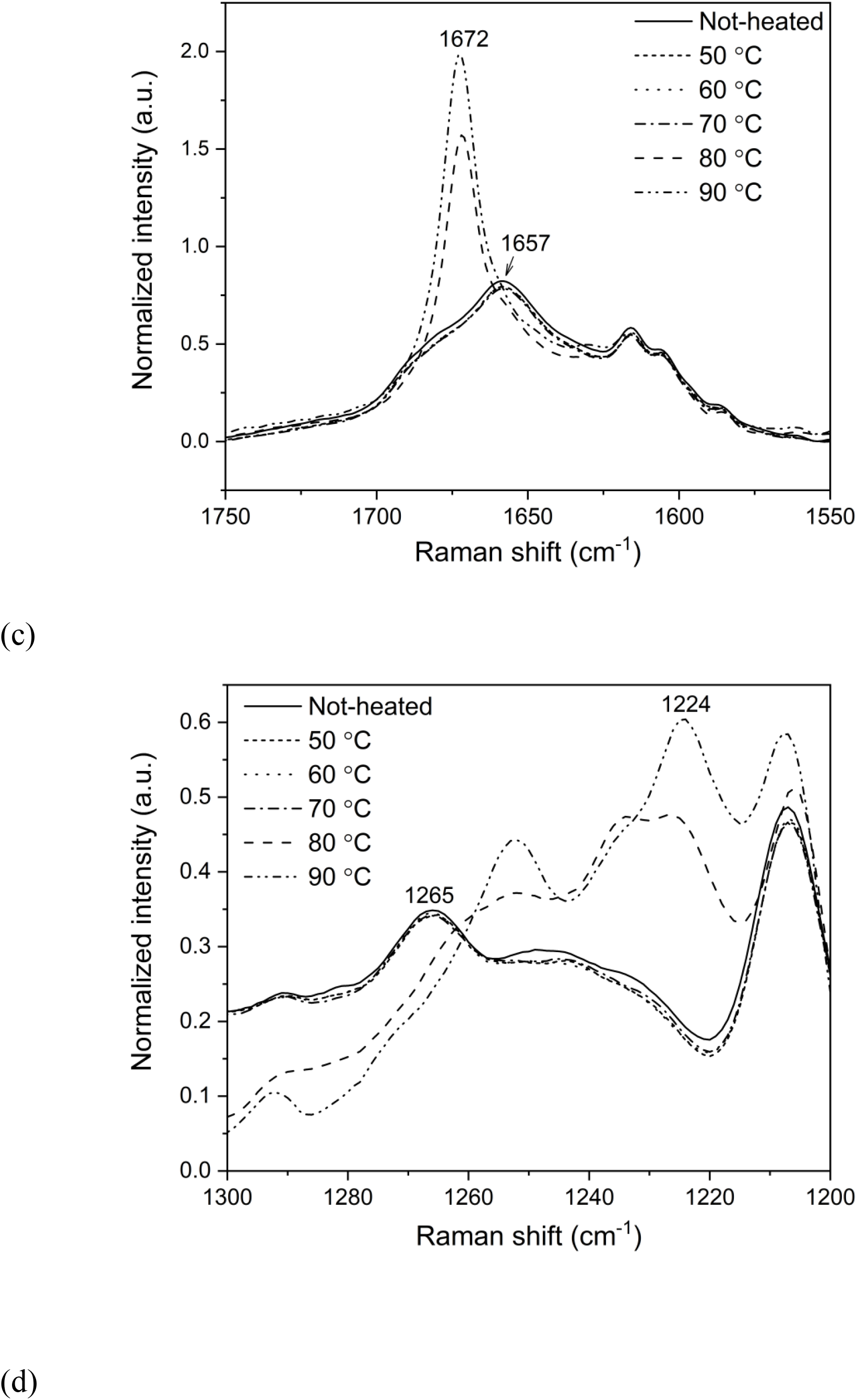
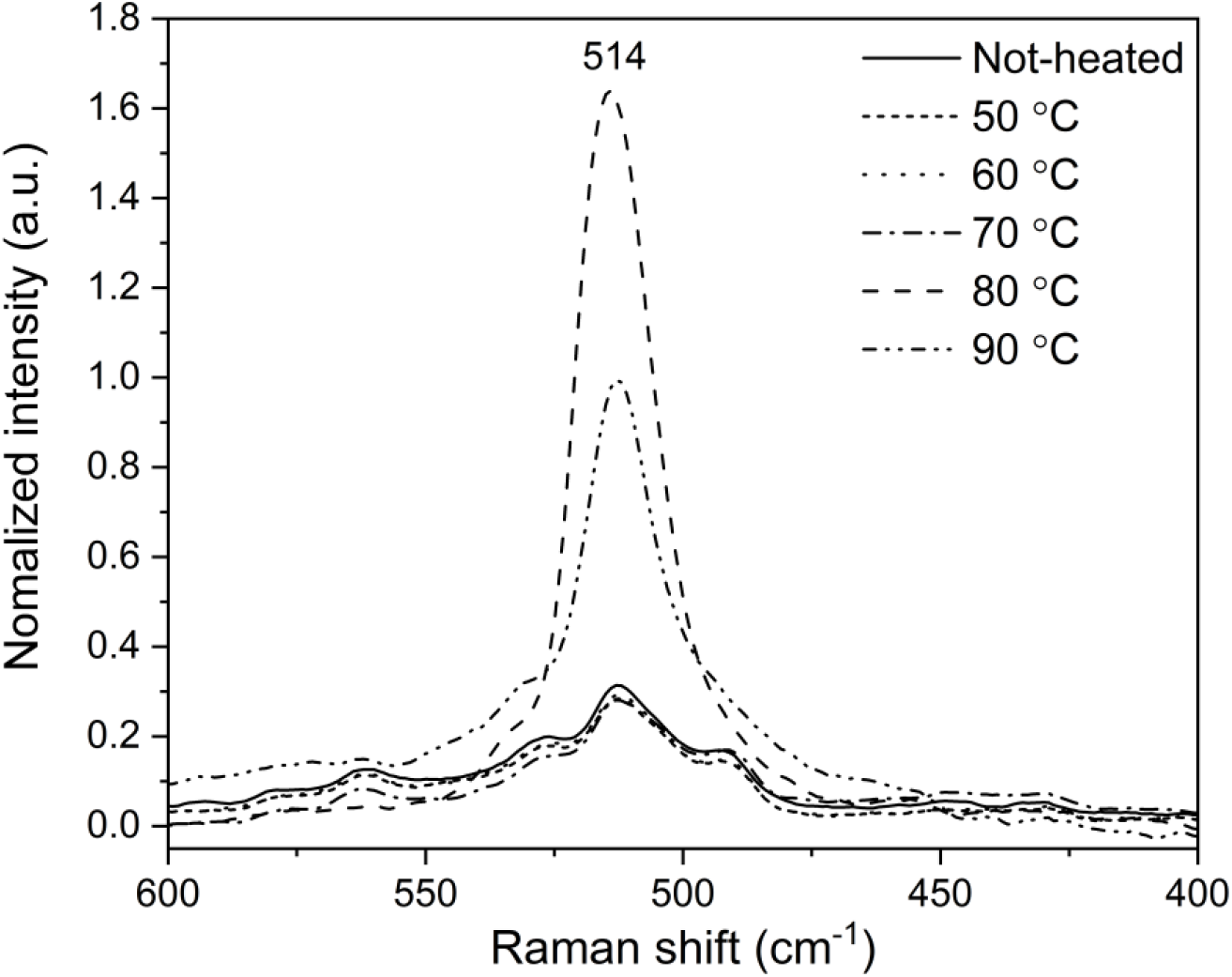
(a) Raman spectra of human insulin solutions in the 1800-300 cm^-1^ wavenumber region that were heated at each temperature for 20 minutes. The magnified spectra of amide I and III regions in (b) 1750-1550 and (c) 1300-1200 cm^-1^ wavenumber regions. (d) Raman peaks in the 600-400 cm^-1^ range due to S-S vibrational mode.

In order to quantitatively evaluate the progress of amyloid formation, the intensity ratio of Raman peaks of the β-sheet (1672 cm^-1^) and α-helix (1657 cm^-1^) in the amide I region defined as *R* (*I*_1672_⁄*I*_1657_) was plotted in Figure 2a. The evidence of amyloid formation is normally obtained using ThT fluorescent dye (43). Alternatively, the band areas of the IR spectra due to the α–helix and β–sheet structures of proteins are also plotted as evidence for amyloid formation (44). In the present study, the structural changes of the protein expressed as temperature-dependent Raman data *R* (*I*_1672_⁄*I*_1657_) showed a similar trend as that of the variation in ThT fluorescence intensity shown in Figure 2b. The present results proved that amyloid fibril formation could be pursued by assessing the ratio of Raman band intensities due to α-helix and β-sheet.

**Figure 2:**
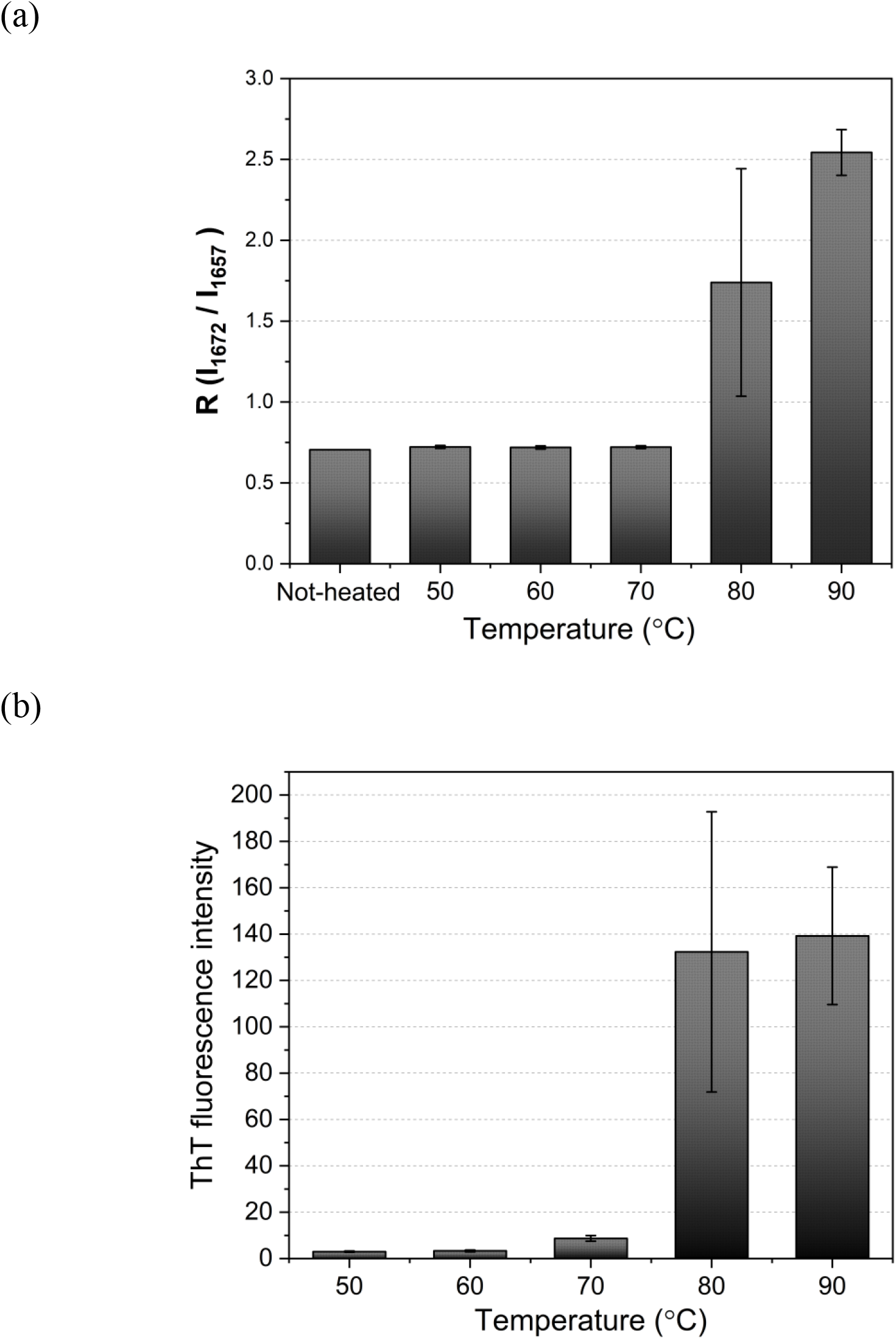
(a) The ratio of peak intensity defined as *R*(*I*_1672_⁄*I*_1657_). (b) ThT fluorescence intensity depending on heating temperature.

The heating time was changed at four different temperatures (60, 70, 80, and 90 ℃). Figure 3a demonstrates the spectral variations in the amide I region (1750-1550 cm^-1^) due to the time course of sample heating at 60 ℃. The appearance of amide I band owing to β-sheet structure (1672 cm^-1^) was observed after 55 minutes of heating. The ratio of Raman peaks *R*(*I*_1672_⁄*I*_1657_) was calculated for each heating temperature (Figure 3b), and the data plots were fitted as a sigmoid curve using the Boltzmann function defined below.

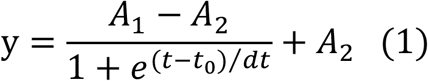

**Figure 3:**
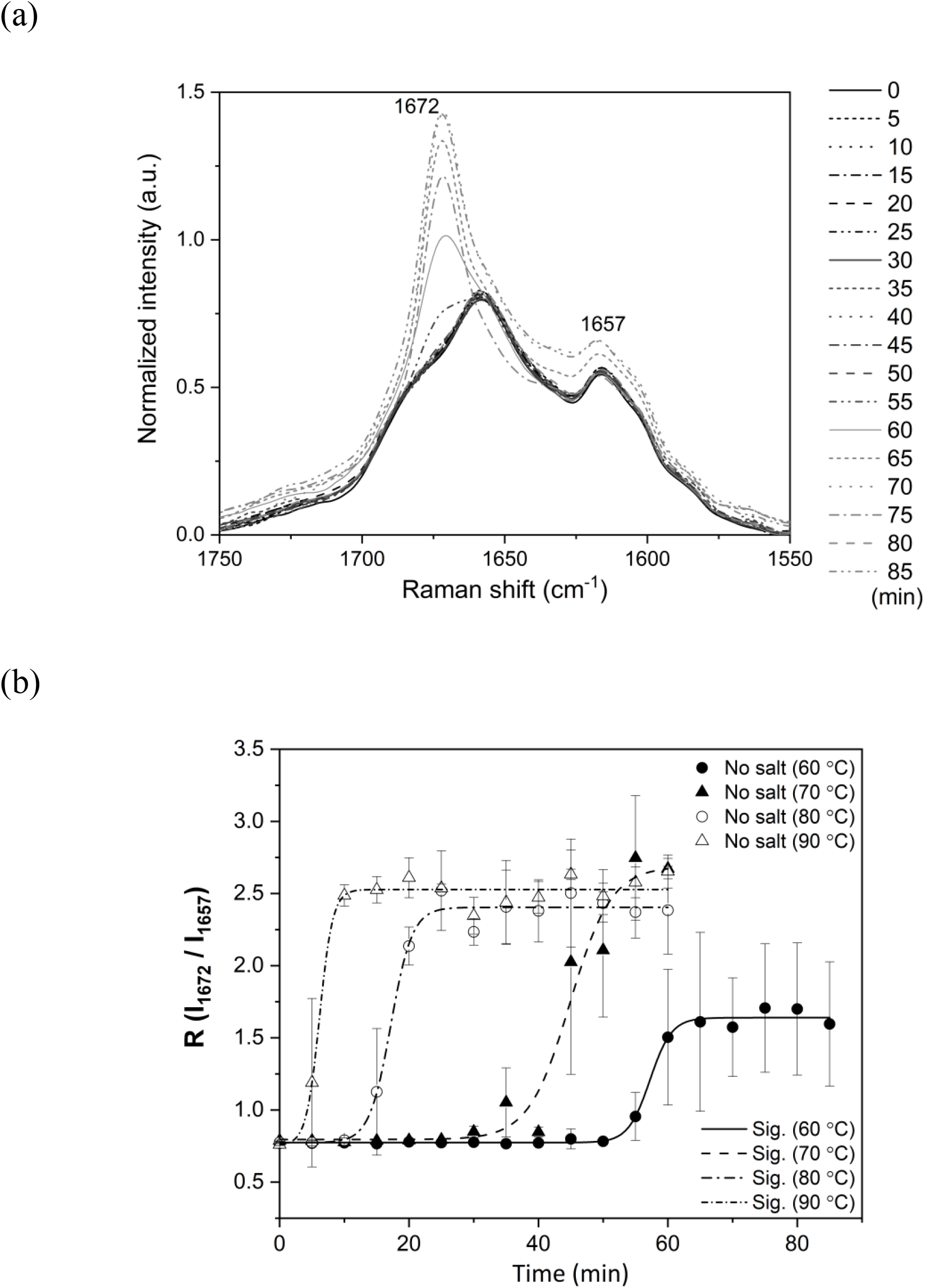
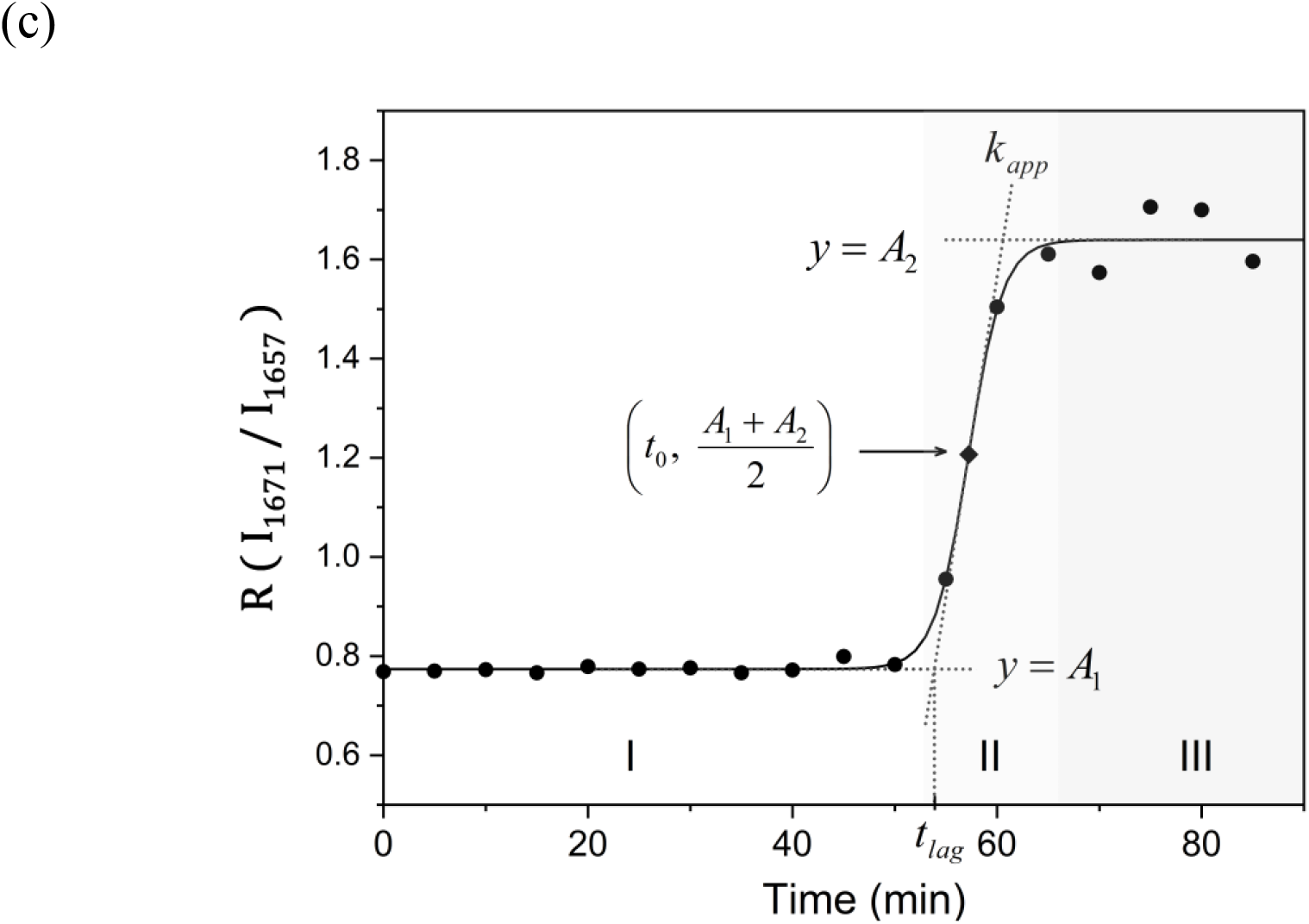
(a) The spectral variation at around amide I region (1750-1550 cm^-1^) due to the time course of sample heating at 60 ℃ without any salts. (b) The plotof the ratios of peak intensities *R*(*I*_1672_⁄*I*_1657_). (c) The definitions of Boltzmann parameters, *k*_*app*_ and *t*_*lag*_.

Here, *A*_1_ and *A*_2_ are y values of an approximate straight line drawn parallel to the y axis representing *R*(*I*_1672_⁄*I*_1657_) (Figure 3c). Amyloid fibril formation consists of three phases: nucleation (I), elongation (II), and equilibrium (III) phases (45, 46). A tangent was drawn at the point *x*_0_ at which the y value was y = (*A*_1_ + *A*_2_)⁄2, and the slope of the tangent was defined as an apparent elongation rate (*k*_*app*_) of amyloid fibrils. Furthermore, the *t* value of the tangent intersection in nucleation and elongation phases was defined as the lag time (*t*_*lag*_) until the beginning of fibrillation; the higher the temperature, the shorter the lag time. The values of *R* (*I*_1672_⁄*I*_1657_) were lower at 60 ℃ than at other temperatures, and this indicated the differences in the proportion of protein structures with a β-sheet depending on the heating temperature.

The time course monitoring for amyloid formation was also observed for samples with the addition of two species of salts (NaCl and Na_2_SO_4_) and their controls. Figure 4a depicts the Raman spectra of insulin samples with and without salts before heating in order to investigate any spectral differences from native insulin caused by salts effects. There were no specific changes to protein structures caused by salt effects except for the small peak at 980 cm^-1^ due to the symmetric stretching vibration of the SO_4_ ^2-^ of the salt (47, 48). Time-dependent spectral variations in the insulin samples with the salts, showing the secondary structural change from α-helix to β-sheet with heating were also observed in the amide I and III wavenumber regions with the formation of amyloid fibrils. Figure 4b and 4c show spectral variations in the 1750-1550 cm^-1^ region depending on the heating time at 60 ℃ with NaCl and Na_2_SO_4_, respectively. With NaCl, the secondary structure started to vary at around 55 minutes and in the case of Na_2_SO_4_, on the contrary, it started to vary at around 40 minutes. Due to the salt effects, the fibril formation seemed to occur early with Na_2_SO_4_ at 60 ℃. The ratios of the two Raman peaks *R*(*I*_1672_⁄*I*_1657_) in Figure 4d and 4e show the time course of the fibrillation process with NaCl and Na_2_SO_4_, respectively. Table 1 summarizes the results about *t*_*lag*_ (± SD) and *k*_*app*_ (± SD) with and without salts at various heating temperatures. The *t*_*lag*_ tended to get shorter and the *k*_*app*_ became higher with the increase of the heating temperature in all the three samples. That is, the amyloid nucleation occurred in a shorter time and the amyloid fibrils were immediately elongated at a higher temperature. This relationship between *t*_*lag*_ and *k*_*app*_ in which the nucleation and elongation of amyloid fibrils were promoted with the increment of heating temperature was consistent with previous studies (15, 49). Furthermore, comparing results with and without salts, the *t*_*lag*_ came earlier, but the elongation speed decreased due to salt effects, especially at lower temperatures. That is, salt injections made it easier for nuclear creation and inhibited the elongation of fibrils. At higher temperature, the variations of *t*_*lag*_ and *k*_*app*_ were negligible and the reaction of fibril formation was severe enough to neglect the salt impact on the speed of amyloid formation. The final values of *R*(*I*_1672_⁄*I*_1657_) in the plateau of the sigmoid curve with Na_2_SO_4_ had variations depending on heating temperature (Figure 3b, 4d, and 4e). The higher proportion of β-sheet structure presented in amyloid fibrils at higher temperature with Na_2_SO_4_.

**Figure 4:**
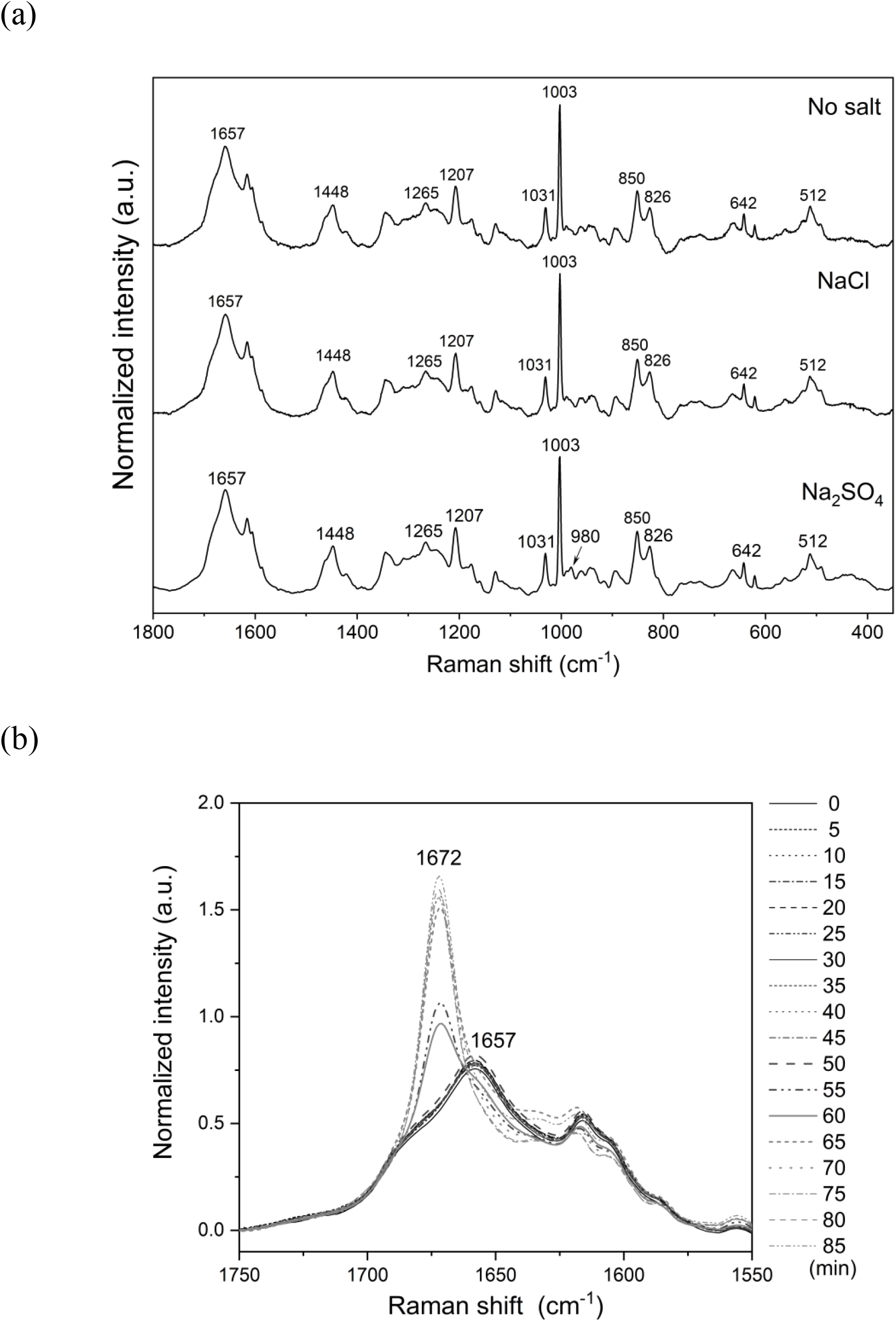
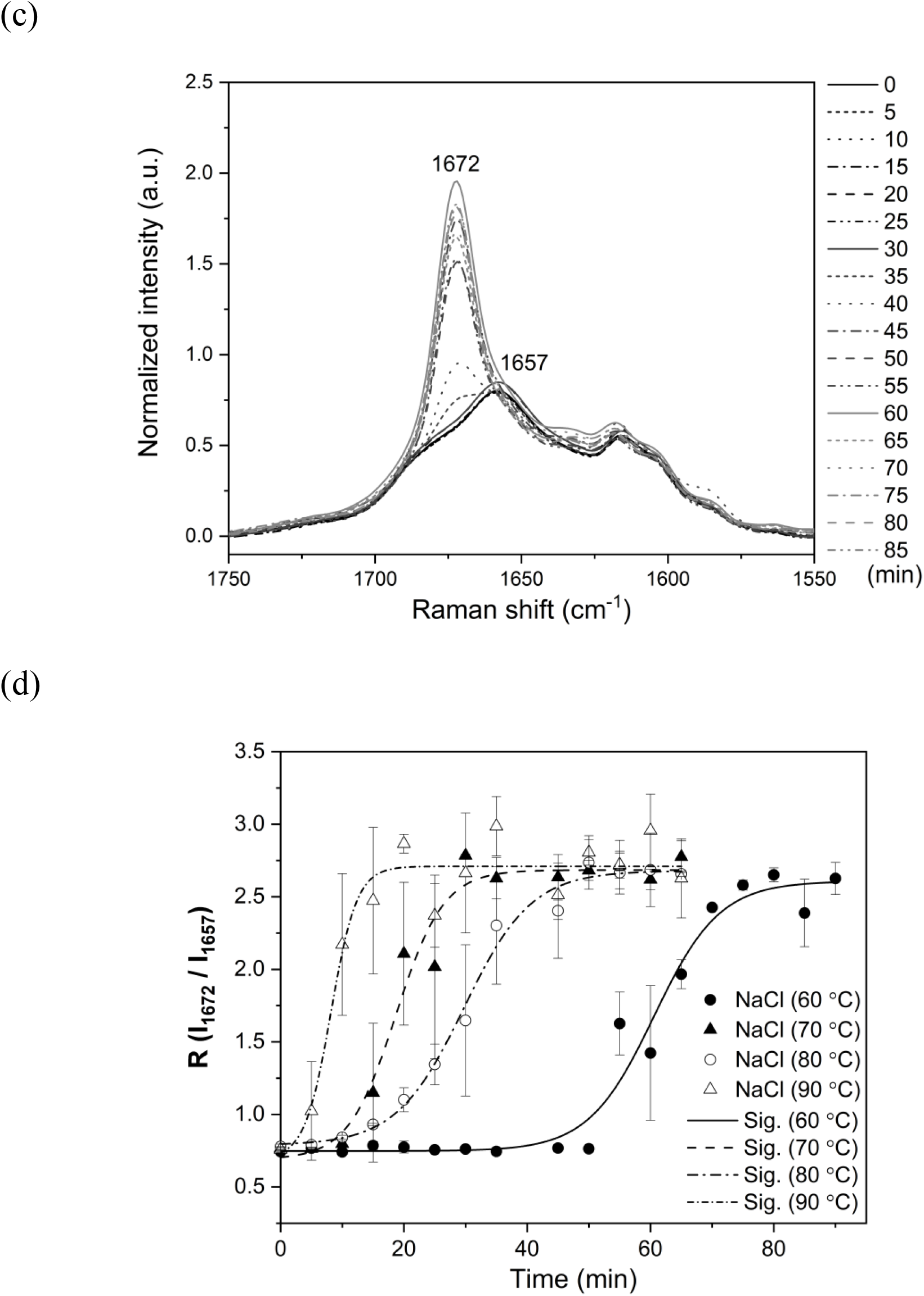
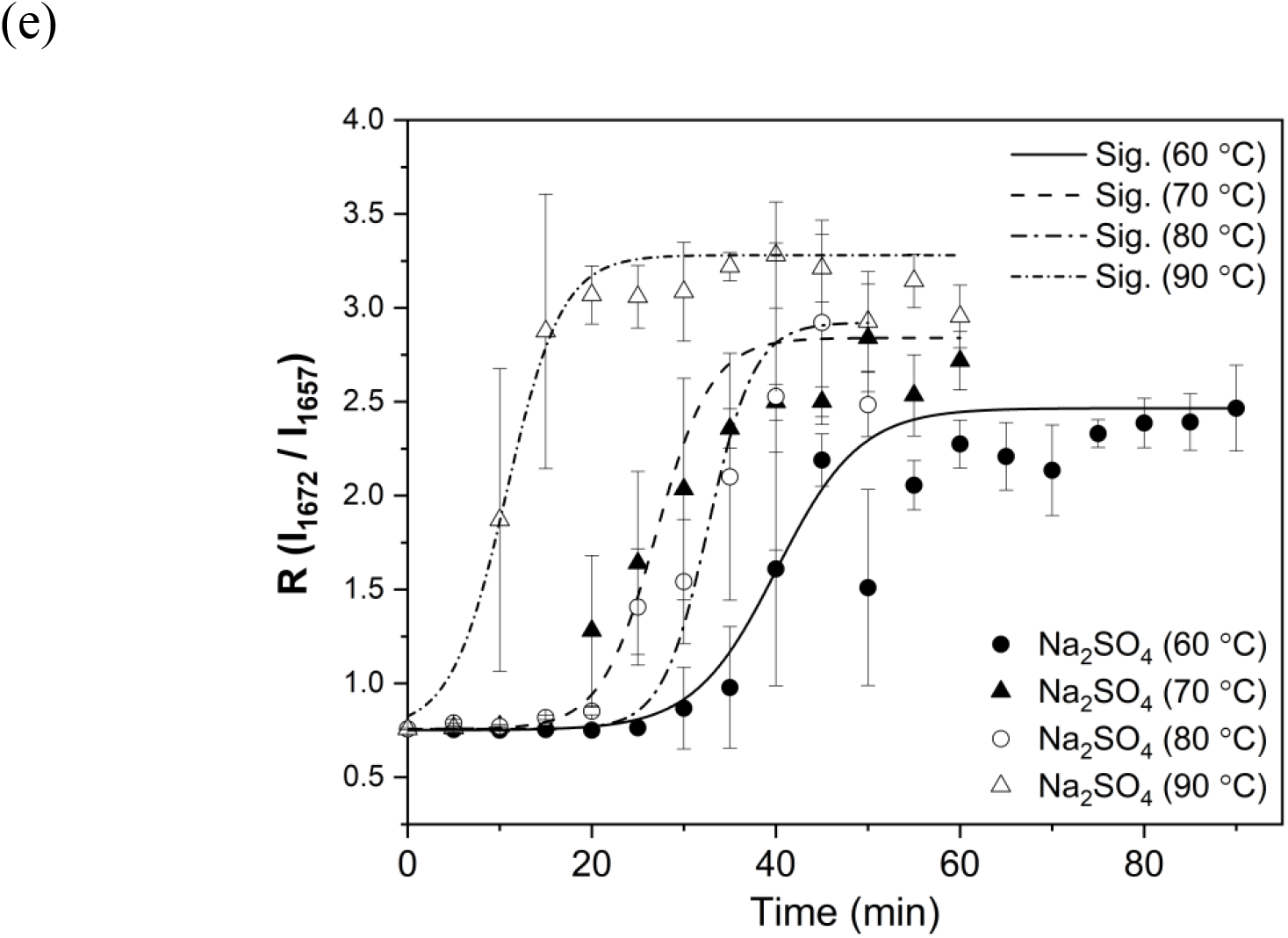
(a) Mean Raman spectra of insulin samples with and without salts before heating. The spectral variations in the 1750-1550 cm^-1^ region depending on heating time at 60 ℃ with (b) NaCl and (c) Na_2_SO_4_. The time-dependent variation of the ratio of Raman peaks *R*(*I*_1671_⁄*I*_1657_) and fitted as sigmoid curves using Boltzmann function with (d) NaCl and (e) Na_2_SO_4_.

**Table 1:** Summary of the results for *t*_*lag*_ and *k*_*app*_.

|  |  | 60°C | 70°C | 80°C | 90°C |
| --- | --- | --- | --- | --- | --- |
| $t_{lag}(\text{min})$ | No salt | $60.6 \pm 0.54$ | $51.2 \pm 2.4$ | $20.5 \pm 0.7$ | $8.2 \pm 1.2$ |
| | NaCl | $71.1 \pm 2.3$ | $26.7 \pm 2.3$ | $39.7 \pm 1.9$ | $12.0 \pm 1.4$ |
| | Na<sub>2</sub>SO<sub>4</sub> | $54.7 \pm 4.8$ | $36.5 \pm 2.1$ | $40.6 \pm 3.2$ | $14.0 \pm 1.1$ |
| $k_{app}(\text{min}^{-1})$<br>($\times 10^{-3}$) | No salt | $12.9 \pm 1.7$ | $14.9 \pm 5.0$ | $24.6 \pm 4.2$ | $43.1 \pm 22.5$ |
| | NaCl | $8.8 \pm 1.7$ | $12.3 \pm 3.2$ | $9.4 \pm 1.6$ | $26.1 \pm 8.2$ |
| | Na<sub>2</sub>SO<sub>4</sub> | $5.9 \pm 1.9$ | $9.1 \pm 1.6$ | $10.4 \pm 3.2$ | $31.7 \pm 8.3$ |

### Polymorphism of amyloid fibrils caused by salt effects

To investigate the polymorphism of amyloid fibrils caused by salt effects, PCA was performed on the dataset of Raman spectra recorded from fibrillated samples in equilibrium phase III at different temperatures. Figures 5a-d are score plots of PCA at heating temperatures of 60, 70, 80, and 90 ℃, respectively. The data were roughly classified into several groups by salt species and the separation tendency became more prominent above 80 ℃. The datasets for 80 and 90 ℃ were divided into two or three groups (Figure 5c and 5d) by the principal component 1 (PC1). The loading plots of PC1 at 80 and 90 ℃ in Figure 5e show some common peaks at 1672, 1235, 850, and 514 cm^-1^, which are characteristic vibrational modes of the β-sheet structure of proteins (1672 and 1235 cm^-1^), the tyrosine ring breathing (850 cm^-1^), and the disulfide bond (514 cm^-1^) (28, 29). This indicates that the Raman peaks at 1672 and 1235 cm^-1^ obtained from amyloid fibrils with salts, especially for Na_2_SO_4_, are stronger at 80 and 90 ℃, and the ratios of the protein structures transitioned to β-sheet are higher under these conditions. It is consistent with the results derived from the values of *R*(*I*_1672_⁄*I*_1657_) in phase III shown in Figure 3b, 4d, and 4e. The striking peak at 514 cm^-1^ was due to disulfide bond with g-g-g conformation (Figure 5e). Thus, the peak is more intense with Na_2_SO_4_ than in the other two cases. In fact, the peak intensity at around 514 cm^-1^ in the averaged spectra of fibrilized samples heated at 90℃ markedly protruded for the Na_2_SO_4_ sample (Figure 5f).

**Figure 5:**
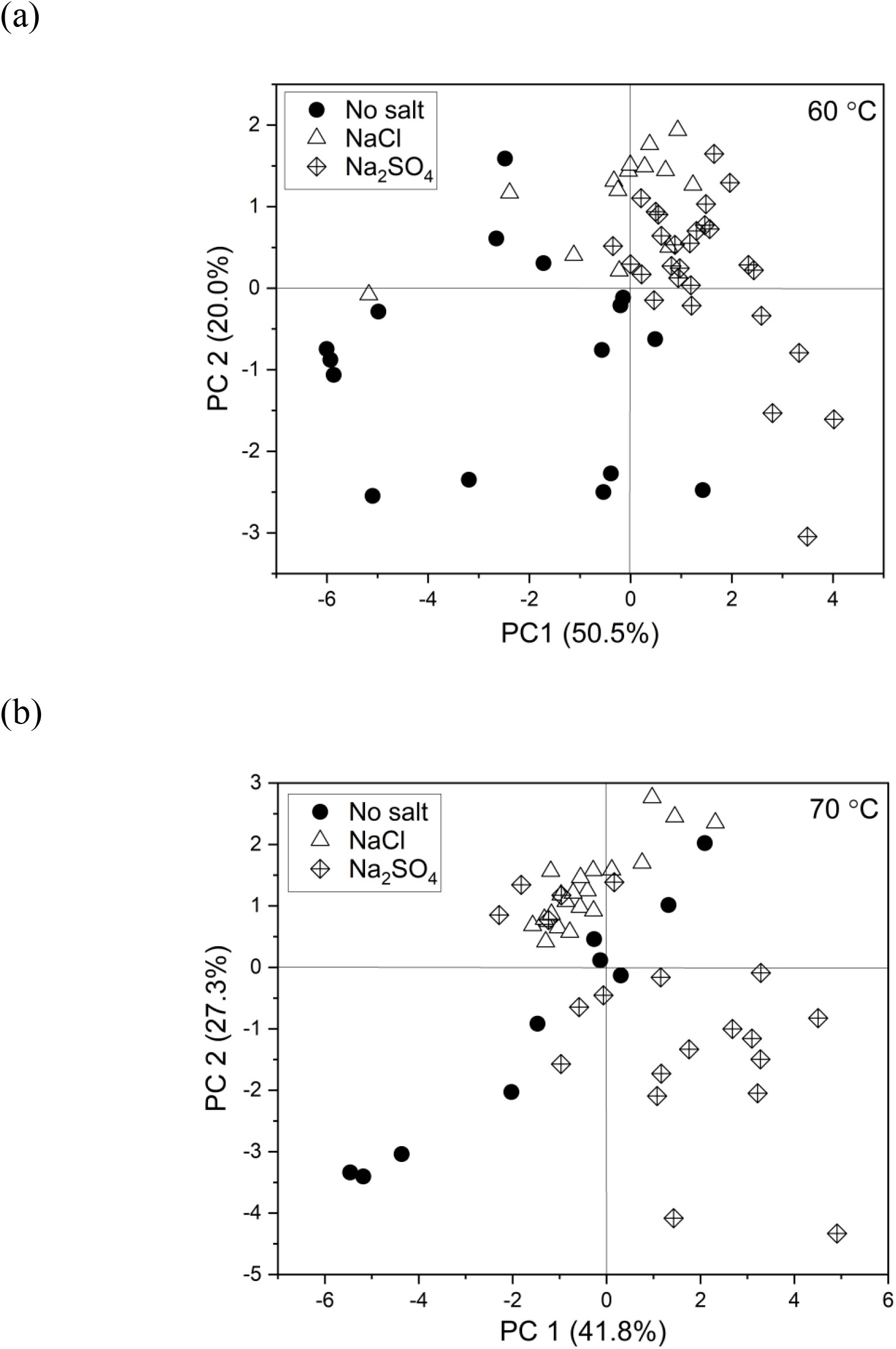
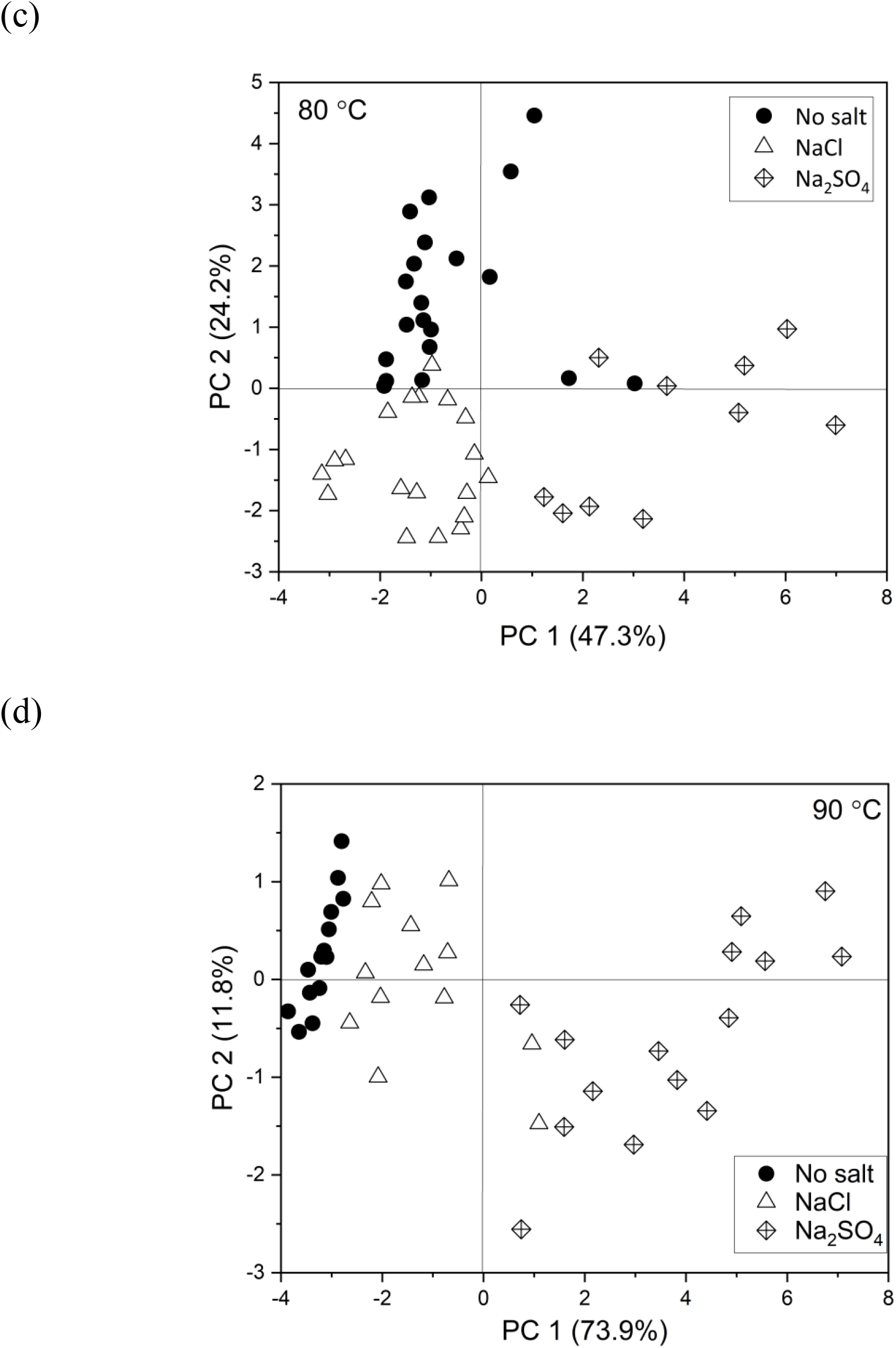
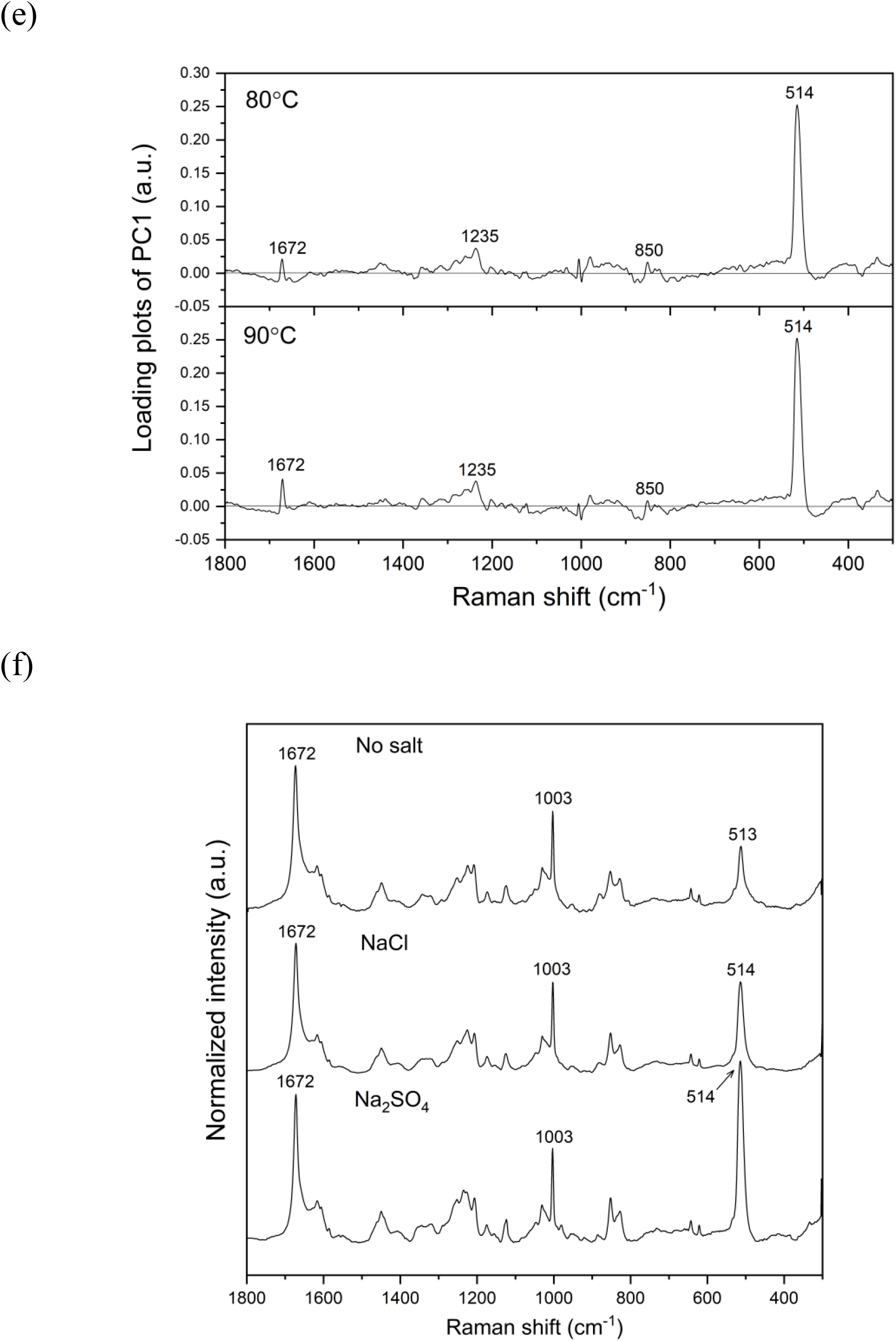
(a)-(d) Score plots of PCA derived from the dataset of different temperatures. (e) Loading plots of PC1 at 80 and 90 ℃. (f) Averaged Raman spectra of fibrillar samples with and without salts heated at 90 ℃.

In addition, the PCA results showed that the Raman peak at 850 cm^-1^ became stronger in the sample with added salt. The ratio of the peak intensities of the tyrosine doublet at around 850 and 826 cm^-1^ [*R* (*I*_850_⁄*I*_826_)] is the barometer of the environment surrounding the hydroxyl group of the tyrosine residue (30). Martel et al. discussed the changes of the ratio of the tyrosine doublet in the structural transition of silk fiber assembly from an α-helix to a β-sheet (50). The ratio of the band intensities varied from almost 4 to 2, and this result was interpreted as the change in the environment surrounding tyrosine residues from being hydrophobic to being more hydrophilic (50). Siamwiz et al. discussed the values of the ratio more generally from the point whether phenolic OH in a tyrosine works as an acceptor or a donor of hydrogen bonds. They estimated that the values lay in the range of 0.9-1.4 when tyrosine residues were on the surface of proteins in aqueous solutions, and the range of 0.3-1.4, in the hydrophobic conditions when tyrosine residues were buried within proteins (30). The values of tyrosine doublet are different depending on samples and the scale of variation, caused by the changes of environment surrounding the tyrosine residues, does not have a fixed value (30). Therefore, just by looking at the values of tyrosine doublet, we cannot conclude anything about the environment of the tyrosine residue. However, the environmental changes of tyrosine residues can be discussed from the variation of the ratio of the tyrosine doublet and consistently understood from the point of the net negative charge of phenolic oxygen within it.

The tyrosine doublet is formed due to the Fermi resonance between the ring breathing vibration (*v*_1_) and the overtone of an out-of-plane ring vibration (*v*_16*a*_) of tyrosine phenyl ring benzenes (30). The ratio of tyrosine doublet can be expressed by the following Equation 2.

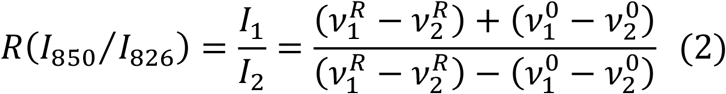

The detailed extraction of Equation 2 is provided in Supporting Information. *I*_*n*_ and 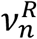 are observed Raman intensities and frequencies of the tyrosine doublet. The subscripts *n* = 1 and *n* = 2 in *I*_*n*_, 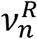, and 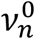 express the ring breathing vibration and the out-of-plane ring breathing vibration of tyrosine benzenes, respectively. By applying experimental data to Equation 2, the separation between 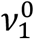 and 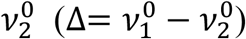 before Fermi resonance can be calculated.

It is well known that the intensity ratio of two band intensities is sensitive to the state of the phenolic hydroxyl group. The net negative charges in the atoms of phenolic oxygen are increased by hydrogen bonds (30). It has been reported that the increment of net negative charge increases the Raman band position of tyrosine doublet with lower frequency (2*v*_16*a*_) to higher wavenumber (30). In that case, the decrement of value Δ enhanced the denominator of Equation 1, and as a result, the ratio of the tyrosine doublet was eventually suppressed.

Table 2 summarizes the observed values of tyrosine doublet *R*(*I*_850_⁄*I*_826_), the observed wavenumber of two tyrosine bands, and the expected Raman shift separation (Δ) between 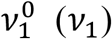 and 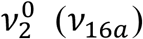 with the SD. Four tyrosines are included in an insulin molecule and each one is in a different condition within the fibrilized samples. Therefore, the ratio of the tyrosine doublet and Δ inevitably seem to have large variations. However, the values before and after heating at 90 ℃ were significantly different among the samples with and without salts. At first, the samples with salts before heating had lower values of Δ and anions might have introduced bigger negative charge of oxygen atoms in tyrosine residue. Collins et al. proposed the model dynamics of salts affecting the hydration water of proteins (51). Anions, such as 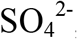, strongly bind molecules of bulk water and strip the hydration water of proteins. As a result of exposed charge of protein, protein molecules lose their stability and easily form aggregation (51). Therefore, the lower values of tyrosine doublet and Δ in the cases with salts were interpreted as stronger electrical interaction in the proteins due to anions. When amyloid fibrils were formed by heating, the Δ tended to become larger in all cases. The polarization of oxygen atoms in tyrosine residues were likely to be reduced by crawling into the hydrophobic environment of amyloids, and the decrease of intensity *I*_2_ seemed to increase the ratio *R*(*I*_850_⁄*I*_826_). Furthermore, the insulin samples with salts had larger values of *R*(*I*_850_⁄*I*_826_) and Δ than those without salts at 90℃. This indicated that the negative charge of phenolic oxygen with salts was relatively small compared to those without salts at higher temperature by being more buried in aggregations. That is, present results comparing the ratio of tyrosine doublet *R*(*I*_850_⁄*I*_826_) and Δ revealed the polymorphisms of amyloid fibrils due to salt effects and heating temperature.

**Table 2:** Summary of the values of tyrosine doublet *R*(*I*_850_⁄*I*_826_), measured frequency of tyrosine bands, and the expected Raman shift separation (Δ) between 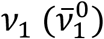 and 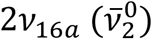 before Fermi resonance. (BH: Before heating)

|  |  | BH | 60°C | 70°C | 80°C | 90°C |
| --- | --- | --- | --- | --- | --- | --- |
| $R(I_{850}/I_{826})$ | No salt | $1.47 \pm 0.05$ | $1.58 \pm 0.20$ | $1.40 \pm 0.18$ | $1.41 \pm 0.18$ | $1.28 \pm 0.13$ |
| | NaCl | $1.29 \pm 0.05$ | $1.35 \pm 0.07$ | $1.54 \pm 0.13$ | $1.35 \pm 0.10$ | $1.48 \pm 0.09$ |
| | Na<sub>2</sub>SO<sub>4</sub> | $1.31 \pm 0.09$ | $1.54 \pm 0.12$ | $1.55 \pm 0.10$ | $1.43 \pm 0.16$ | $1.47 \pm 0.11$ |
| The<br>wavenumber of<br>Raman peaks in<br>Tyr. doublet<br>(cm<sup>-1</sup>) | No salt | $851.0 \pm 0.5$ | $852.1 \pm 0.9$ | $851.9 \pm 0.3$ | $853.0 \pm 0.7$ | $853.4 \pm 1.5$ |
| | | $826.8 \pm 0.6$ | $827.5 \pm 1.1$ | $827.4 \pm 0.8$ | $827.8 \pm 1.0$ | $828.6 \pm 0.9$ |
| | NaCl | $851.0 \pm 0.5$ | $852.5 \pm 0.5$ | $853.0 \pm 0.8$ | $852.6 \pm 0.5$ | $852.4 \pm 0.6$ |
| | | $827.6 \pm 0.7$ | $829.2 \pm 0.8$ | $828.8 \pm 1.0$ | $827.2 \pm 0.8$ | $826.8 \pm 0.7$ |
| | Na<sub>2</sub>SO<sub>4</sub> | $851.5 \pm 0.5$ | $853.0 \pm 0.8$ | $853.3 \pm 0.5$ | $852.5 \pm 0.6$ | $852.3 \pm 0.6$ |
| | | $827.9 \pm 0.5$ | $829.3 \pm 1.1$ | $829.0 \pm 0.5$ | $827.7 \pm 1.2$ | $827.6 \pm 1.2$ |
| $\Delta$ (cm<sup>-1</sup>) | No salt | $4.59 \pm 0.41$ | $5.50 \pm 1.54$ | $5.58 \pm 0.69$ | $4.25 \pm 1.56$ | $3.04 \pm 1.31$ |
| | NaCl | $2.98 \pm 0.46$ | $3.50 \pm 0.57$ | $5.17 \pm 1.03$ | $3.83 \pm 0.89$ | $4.99 \pm 0.78$ |
| | Na<sub>2</sub>SO<sub>4</sub> | $3.16 \pm 0.78$ | $5.07 \pm 0.94$ | $5.24 \pm 0.80$ | $4.40 \pm 1.38$ | $4.68 \pm 0.90$ |

### Polymorphism of amyloid fibrils caused by heating

PCA was applied to the dataset of each salt species to investigate the effect of temperature on the polymorphism of amyloid fibrils. Figures 6a-c show score plots of PCA for each media. The dataset without salts heated at 90 ℃ was clearly grouped by the PC3 component. Also, the samples with Na_2_SO_4_ were classified into two groups as 60, 70 ℃ and 80, 90 ℃ by PC1 and PC3. The Raman dataset of the samples with NaCl, on the other hand, did not have a special pattern for classification depending on heating temperatures. Figures 6d and 6e exhibit the loading plots of PC1 and PC3 without salts and with Na_2_SO_4_, respectively. Figure 6d shows striking peaks at 514 and 1672 cm^-1^ in the plus direction in PC1 and at 516 cm^-1^ in the minus direction in PC3. The peaks arising from the β-sheet structure and disulfide bonds also appeared at 511 and 1672 cm^-1^ in PC1 and 527 and 1676 cm^-1^ in PC3 in the plus direction, and at 508 cm^-1^ in PC3 in the minus direction in the case with Na_2_SO_4_ (Figure 6e). The peaks at 1672 and 1676 cm^-1^ of PC3 in the plus direction in Figure 6d and 6e indicate that the ratios of β-sheet structure were higher at 80 and 90 ℃ in both cases. Furthermore, the combination of disulfide bonds between 514 and 516 cm^-1^ in opposite directions (Figure 6d) showed a peak shift of the disulfide band to the lower wavenumber at higher temperatures without salts. The peaks at 508 and 527 cm^-1^ in Figure 6e indicate the decrement of the band at around 527 cm^-1^ at higher heating temperatures in addition to the peak shift of disulfide band at lower wavenumber with Na_2_SO_4_. Figures 6f and 6g depict Raman peaks in the 540-500 cm^-1^ wavenumber region without salts and with Na_2_SO_4_, respectively, which are due to disulfide bonds with different conformations; the band at around 510 cm^-1^ comes from the g-g-g conformation and that at 525 cm^-1^ comes from the g-g-t conformation (22, 28, 32, 42). Without salts, the band intensities due to g-g-g conformation (∼510 cm^-1^) decreased but ones due to g-g-t (∼525 cm^-1^) alternatively increased with the increment of heating temperature (Figure 6f). That is, the conformations of disulfide bonds were partially transformed from g-g-g to g-g-t without salts at a higher heating temperature. With Na_2_SO_4_ on the other hand, the band intensities coming from g-g-g conformation were enhanced with the decrease of the peaks due to g-g-t at around 530 cm^-1^ (Figure 6g). The conformations of disulfide bonds were partially changed from g-g-t to g-g-g with Na_2_SO_4_. Krouski et al. reported that the predominant conformation of disulfide bonds in insulin fibrils was g-g-g (32). The present results also proved the dominant conformation in disulfide bonds was g-g-g. However, slight conformational differences were caused by salt effects and heating temperatures, which could be successfully detected by the variation of band ratio due to different disulfide bond conformations.

**Figure 6:**
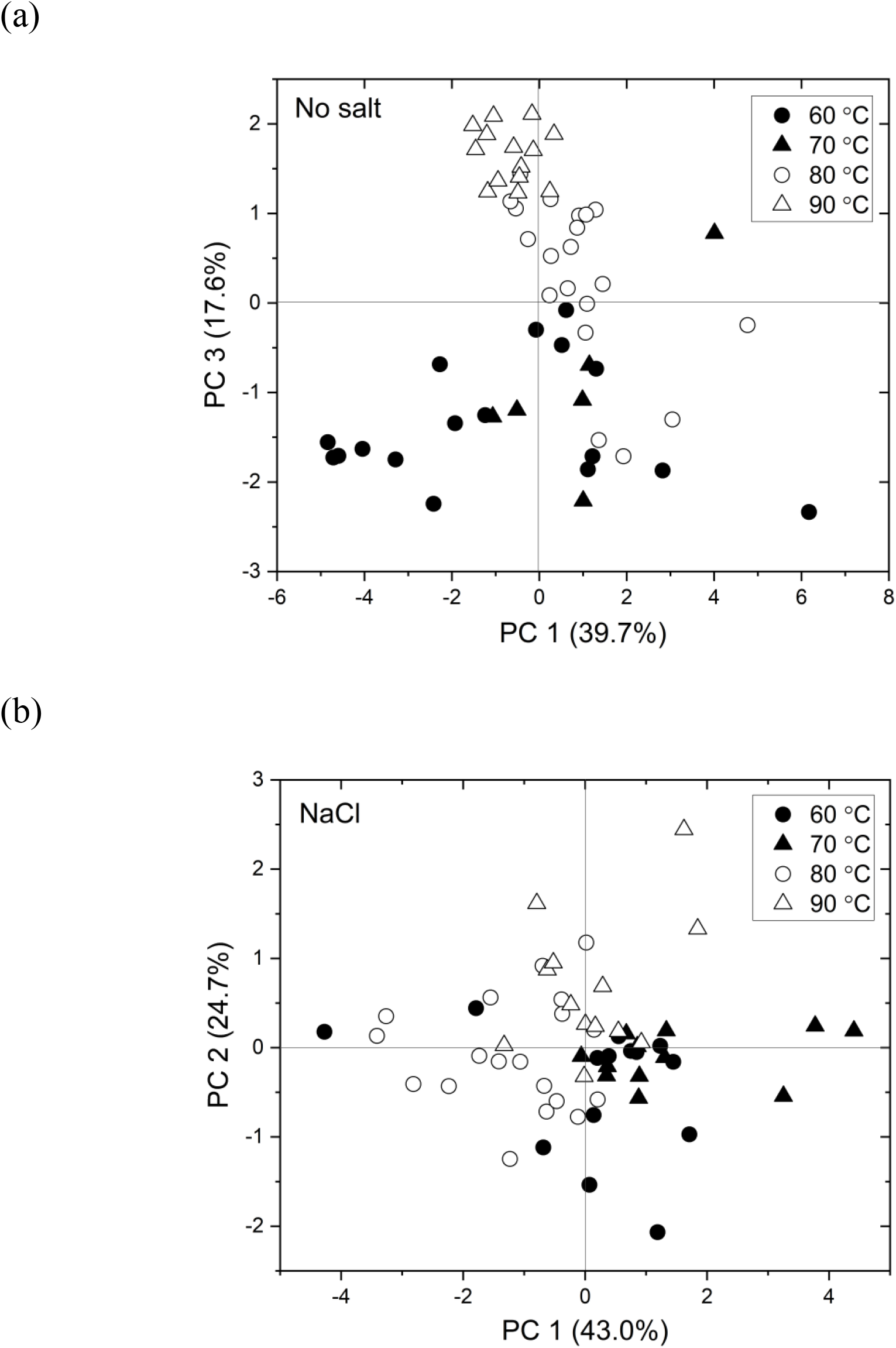
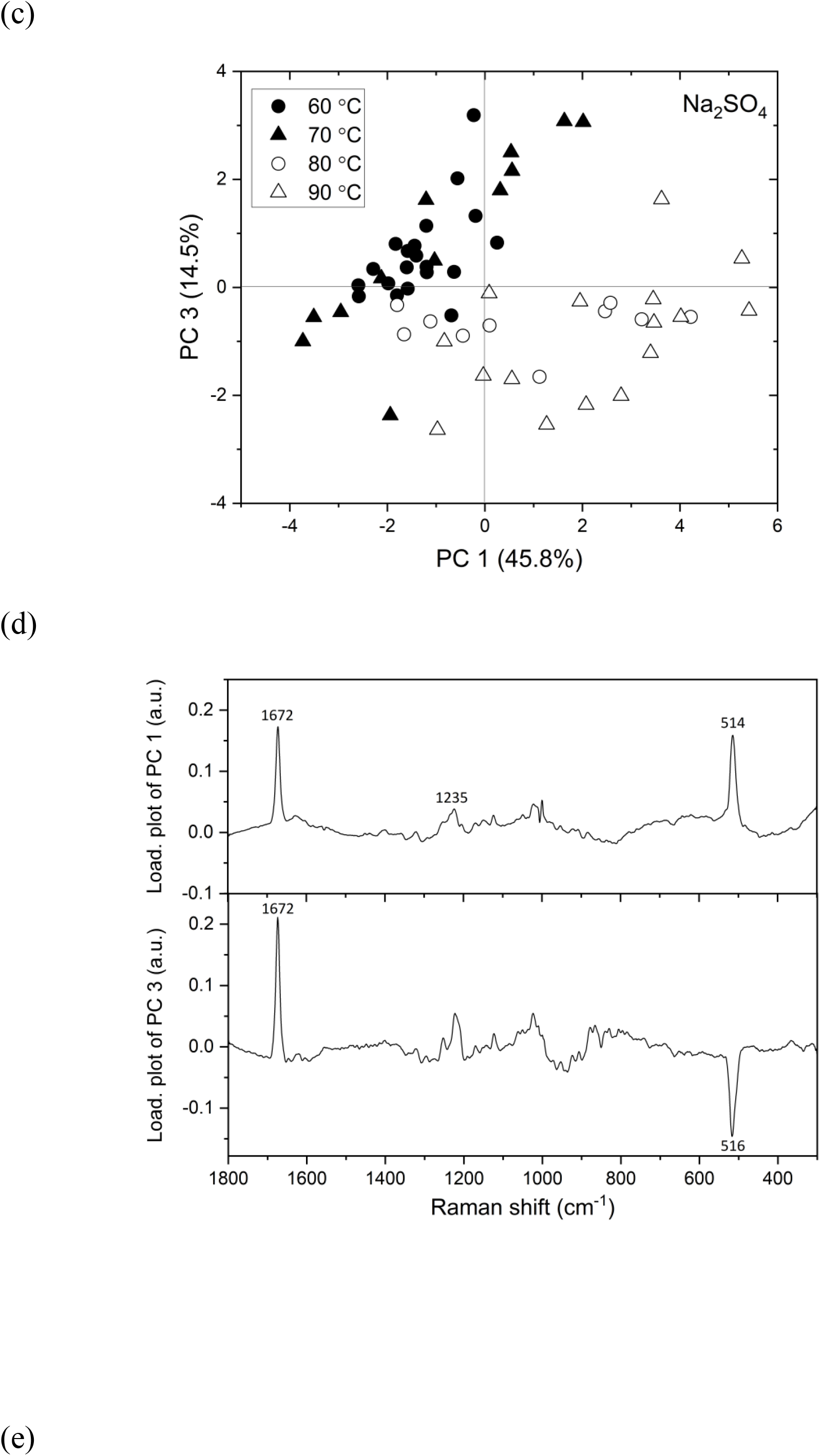
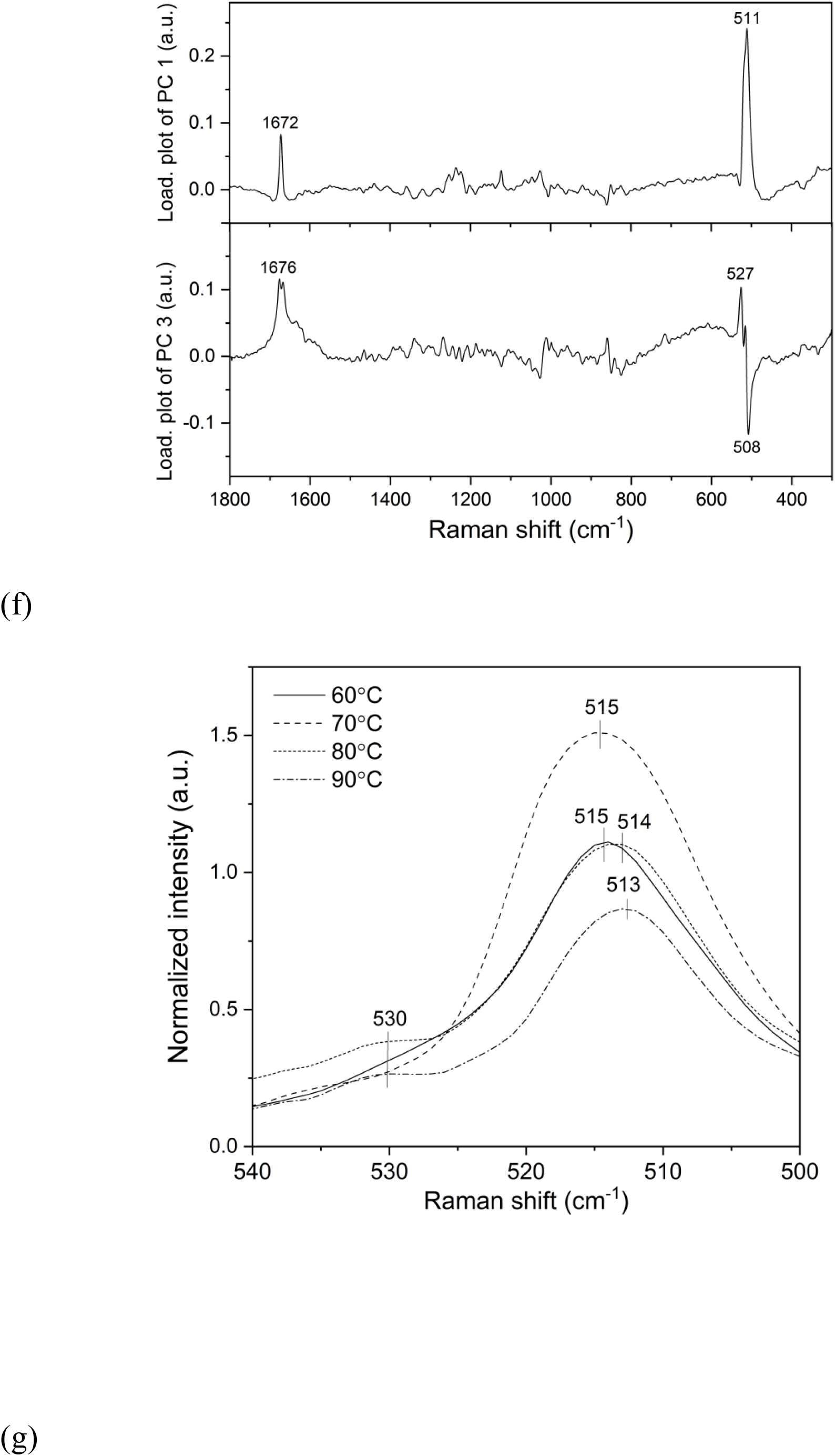
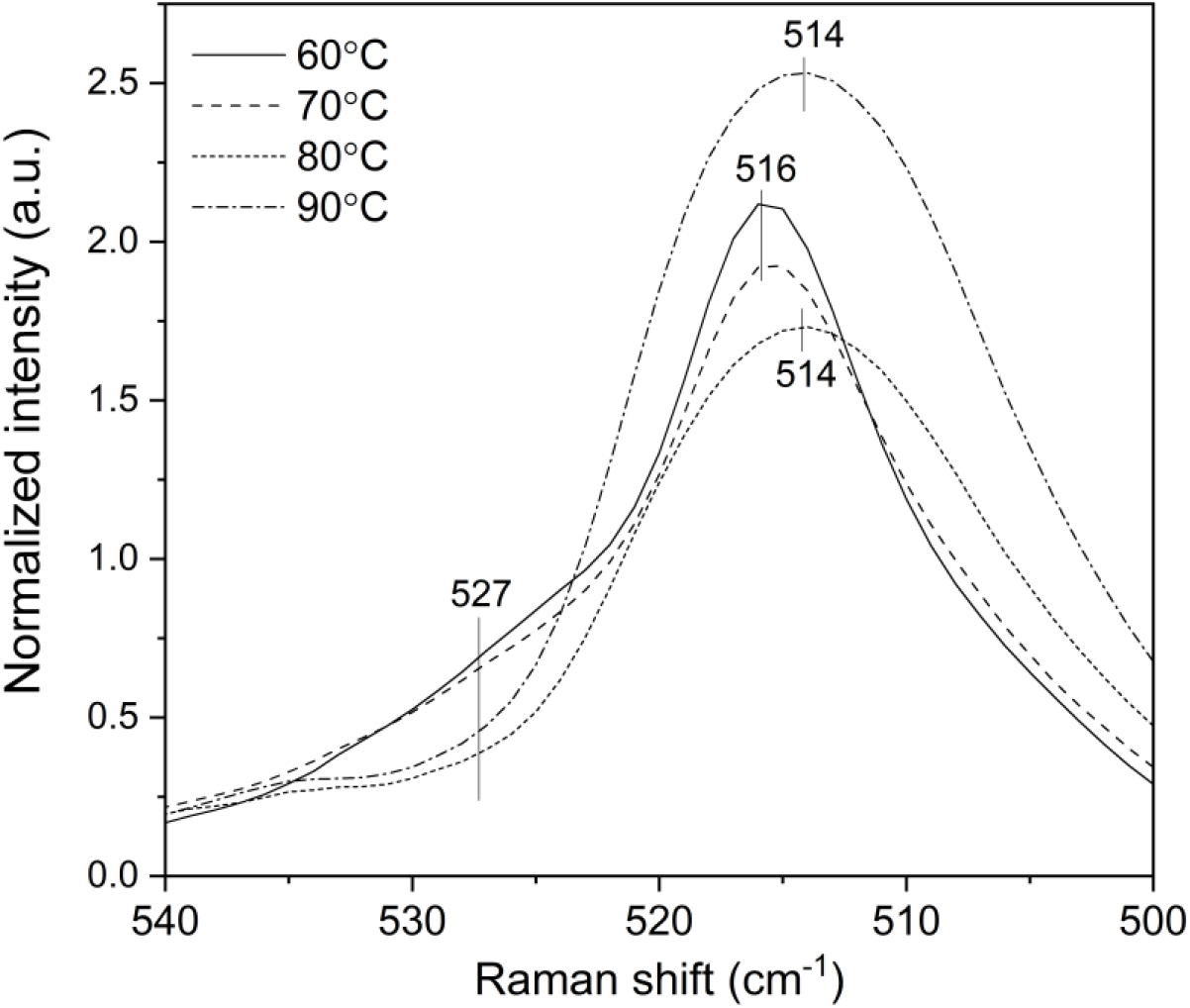
(a)-(c) Score plots of PCA derived from the dataset depending on salt species. The loading plots of PC1 and PC3 (d) without salts and (e) with Na2SO4. The averaged Raman spectra in the 540-500 cm^-1^ wavenumber region (f) without any salts and (g) with Na_2_SO_4_.

### Raman imaging

Raman images were constructed for three kinds of insulin fibrillar aggregates, with and without salts after 30 minutes of heating at 90 ℃. Figures 7 displays (a) visible images of insulin aggregation, and (b)-(e) Raman images developed by plotting the intensities of some notable bands. Figure 7a shows spherical aggregates of several tens of micrometers in size, which are conceived to be spherulites, a kind of hierarchical structure of amyloid fibrils that tend to form at relatively higher concentrations of insulin (52). Figure 7b was constructed using Raman band intensity at 1003 cm^-1^ due to the ring breathing mode of phenylalanine residues. It depicts the distribution of insulin concentration. The spherulite-like aggregates were clearly visualized in agreement with the images shown in Figure 7a, and they were confirmed to be made of insulin. In the case without any salts (i) and with Na_2_SO_4_ (iii), the contour parts of aggregations were more highlighted than the central part. This indicates that the contour of the aggregation was bulky, and that it had concave forms. The Raman images in Figure 7c show the distribution of band intensity at 1672 cm^-1^ due to β-sheet structure normalized by 1003 cm^-1^ internal standard at each point in two dimensions. The aggregates had a higher ratio of β-sheet structure in the order of Na_2_SO_4_ > NaCl > without salts. The Raman images relating to the distribution of β-sheet structure are consistent with those of point mode measurements shown in Figure 5e by PCA. The images in Figure 7d were obtained based on the normalized band intensity at 514 cm^-1^ due to the stretching mode of disulfide bonds with g-g-g conformation. The most interesting pattern was shown in the image with Na_2_SO_4_. The radial structure highlighted inside the aggregate can be seen as a spherulite. Furthermore, inner part of the aggregation has higher intensities compared with those with NaCl and without salts. This is also consistent with the results obtained by point mode measurements described in Figure 5f. Moreover, the disulfide conformations on the surface of the aggregate structure were revealed to be different from those inside the aggregate in both cases, with NaCl and without salts. The images constructed by the ratio of tyrosine doublet in Figure 7e illustrate the environmental variation of tyrosine residues due to addition of salt by constructing the intensity ratio of tyrosine doublet *R*(*I*_850_⁄*I*_826_). The results showed that aggregates had higher values of the ratio, especially with Na_2_SO_4_. The images indicated that the tyrosine was under more hydrophobic conditions by being buried in the aggregates especially with Na_2_SO_4_. Figures 7d and 7e show the higher order structural differences caused by salt effects.

**Figure 7:**
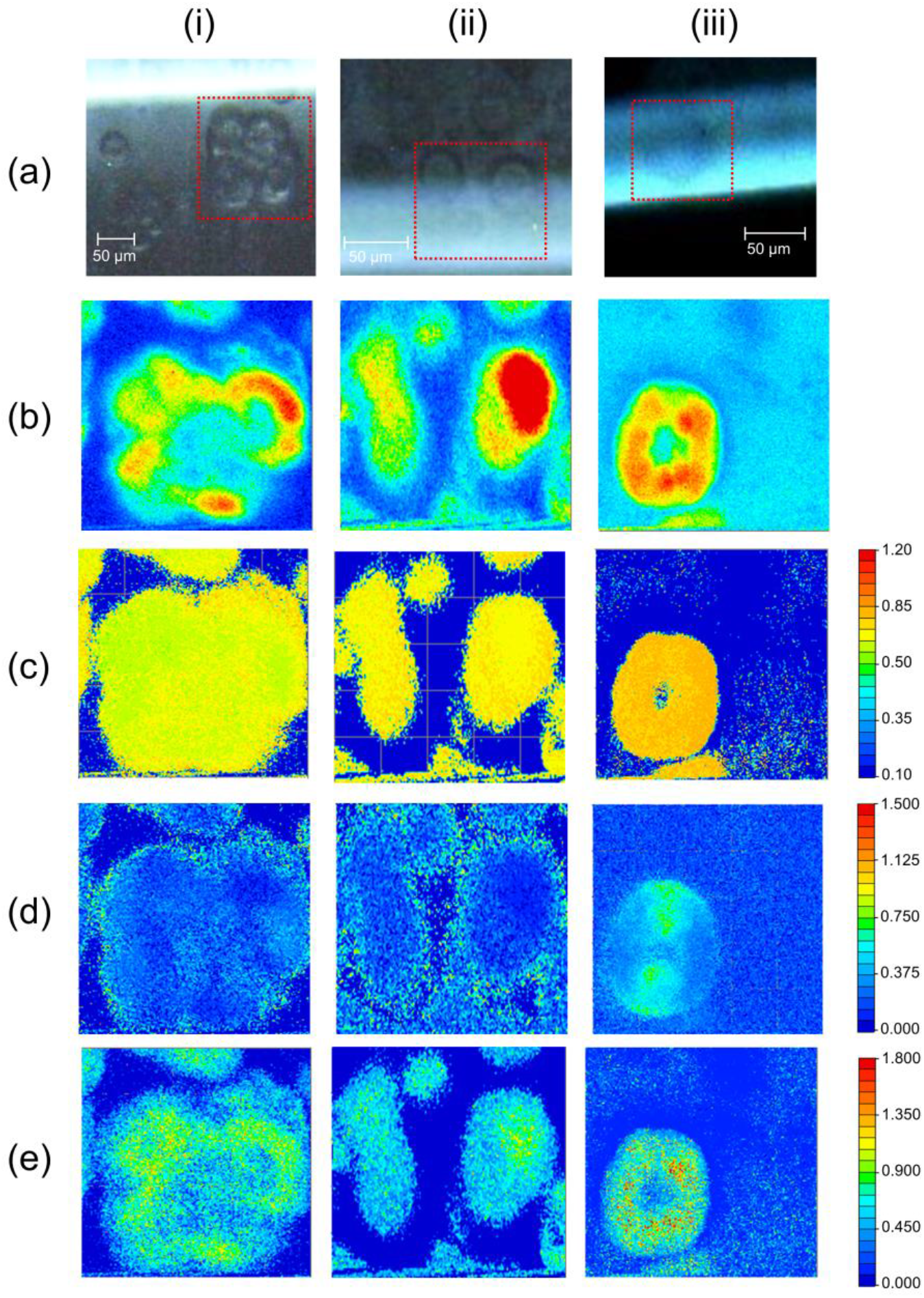
(a) Visible images of insulin aggregation (i) without salts, with (ii) NaCl and (iii) with Na_2_SO_4_. Raman images constructed by plotting Raman band intensities at (b) 1004 cm^-1^, normalized Raman band intensities by a 1004 cm^-1^ internal standard at (c) 1672 cm^-1^ and (d) 514 cm^-1^, and (c) the ratio of tyrosine doublet R(*I*_855_⁄*I*_830_).

**Figure 8:**
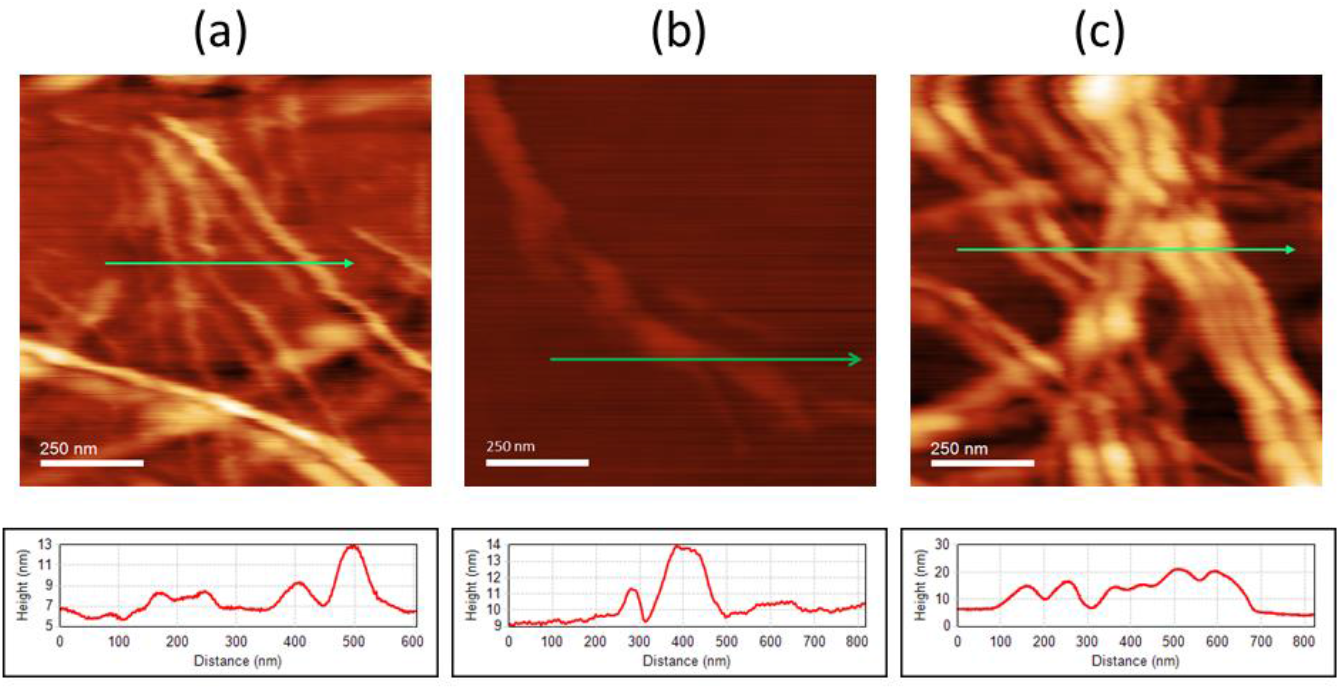
AFM images of amyloid fibrils (a) without salts, (b) with NaCl, and (c) with Na_2_SO_4_.

## Conclusion

The amyloid fibril formation of insulin was investigated *in situ*, focusing on the polymorphism caused by heating temperature and salt effects, using Raman spectroscopy and imaging. The Raman marker bands used to distinguish between polymorphisms of amyloid fibrils were explored by multivariate analysis and three indices were extracted: the relative proportion of β-sheet structure, the intensity ratio of the tyrosine doublet, and the peak intensity due to disulfide bond. First, the relative ratio of the β-sheet structure was revealed to be higher in the case with salts, especially at a higher temperature and with Na_2_SO_4_. The variation of the value *R*(*I*_1672_⁄*I*_1657_) in the time course of fibrillation clearly showed that the lag time until nucleation and elongation speed reached earlier with the increase of heating temperature. Moreover, by adding salts, the lag time was shortened and the elongation speed was lowered compared to those without salts. Second, the intensity ratios of the tyrosine doublet *R*(*I*_850_⁄*I*_826_) changed only by adding salts, before heating. Tyrosine residues in the salt solution were indicated to be surrounded by strong electric interactions. The hydrogen bond network in bulk water caused by salt effects was likely to rob the hydration water from proteins and to induce protein misfolding at earlier stages. Furthermore, the higher values of *R*(*I*_850_⁄*I*_826_) with salts than those without salts at higher temperature were interpreted as tyrosine residues being buried into proteins in hydrophobic aggregations. Lastly, amyloid fibrils with Na_2_SO_4_ media had a higher rate of g-g-g conformation, and in the case without salts, a partially a g-g-t conformation at a higher temperature. Using these indices, polymorphisms of amyloid fibrils were successfully visualized by Raman imaging. Especially, the higher order structural differences were made clear by factors of tyrosine doublet and disulfide bonds. It was very interesting that the surface and inner parts of the aggregates were revealed to have different disulfide conformations, and the ones with Na_2_SO_4_ added were seen to have a spherulite. This work showed how Raman imaging can be interpreted about hydrogen bonds of tyrosine residues based on the detailed discussion about the tyrosine doublet. The present study shows possible applications of Raman imaging to *in situ* diagnosis of amyloid-related diseases.

## Author Contributions

E.C., M.I., and Y.O. designed the study. M.I. and K.M. performed the research. M.I., E.C., and Y.O. wrote the manuscript.

## Acknowledgements

The study was supported by MEXT Leading Initiative for Excellent Young Researchers (M.I.).

